# *Ythdf2* ablation protects aged retina from RGC dendrite shrinking and visual decline

**DOI:** 10.1101/2023.01.05.522855

**Authors:** Fugui Niu, Gaoxin Long, Jian Zhang, Yuanchu She, Jun Yu, Sheng-Jian Ji

## Abstract

Aging-related retinal degeneration and vision loss have been severely affecting the elder worldwide. Previously we showed that the m^6^A reader YTHDF2 is a negative regulator for dendrite development and maintenance of retinal ganglion cells (RGC) in mice ***(Niu et al. 2022)***. Here, we show that conditional ablation of *Ythdf2* protects retina from RGC dendrite shrinking and vision loss in the aged mice. Further, we identify *Hspa12a* and *Islr2* as the YTHDF2 target mRNAs mediating these effects. Together our results indicate that m^6^A modification regulates retinal degeneration caused by aging, which might provide therapeutical potentials for developing new treatment approaches against aging-related vision loss.

## Introduction

Vision loss and blindness in the elder are affecting hundreds of millions of people worldwide, which needs to be addressed as a public health issue with the aging global population ***(Flaxman et al. 2017; Blindness et al. 2021)***. Vision loss in old patients is mostly attributed to aging-related macular degeneration, glaucoma, cataracts, and ocular complication of diabetes mellitus ***(Pelletier et al. 2016)***. Retina, as the fundamental structural tissue to encode and transmit visual signals into the brain, is organized by diverse cell types mediating the signal transduction cooperatively ***(Masland 2012)***. The degenerations in aging retina are associated with such diseases as progressive degeneration of photoreceptors in aging-related macular degeneration and retinal ganglion cells (RGCs) degeneration in glaucoma ***(Weinreb et al. 2014; Fleckenstein et al. 2021)***. In addition, disease-free vision decline is also relevant to structural and physiological changes in retina, including RGC dendrite shrinking, retinal pigment epithelium degeneration, and photoreceptor dysfunction ***(Spear 1993; Jackson et al. 2002; Samuel et al. 2011; Owsley 2016; Datta et al. 2017; Esquiva et al. 2017)***.

Previously we discovered that the m^6^A reader YTHDF2 negatively regulates dendrite development and maintenance of RGCs ***(Niu et al. 2022)***. The expansion of RGC dendrite arbors and more synapses in inner plexiform layer after conditional knockout (cKO) of *Ythdf2* in retina modestly improve the visual acuity of mice in an optomotor assay ***(Niu et al. 2022)***. In the glaucoma models, the m^6^A writers METTL3 and WTAP, and its reader YTHDF2, are upregulated, and loss-of-function of YTHDF2 has a neuroprotective role ***(Qu et al. 2021; Niu et al. 2022)***. Besides, m^6^A modification and METTL3 expression are upregulated under hypoxic and diabetic stress, which governs retinal angiogenesis and pericyte dysfunction of retinal vascular complication ***(Yao et al. 2020; Suo et al. 2022)***. However, it remains unknown whether m^6^A modification and its reader YTHDF2 regulate the degeneration of RGCs in the aged retinas.

Here, we utilized the aged *Ythdf2* cKO mice to analyze the dendrite morphology of RGCs. With the optomotor assay, we detected an improved visual acuity in the aged *Ythdf2* cKO mice. Finally, we identified two target mRNAs which potentially prevent RGCs from degeneration with aging. Therefore, our study indicates that YTHDF2 protects retina from aging-related RGC dendrite shrinking and visual decline, which provides an effective strategy for blocking vision loss in the aged retina.

## Results

### *Ythdf2* deletion increases the dendrite area and branching of RGCs in the aged retinas

The dendrite arbor is the primary information receptive site in a neuron. The geometrical structure of its dendritic arbor determines the receiving region of input, synaptic density, numerous presynaptic partners, and certain physiological properties ***(Lefebvre et al. 2015)***. Rather than neuronal loss, alterations in the dendrite arbors, axonal collaterals, and synaptic density are anatomically detectable in the aged brains ***(Koch et al. 2021)***. In the previous study, we identified the m^6^A reader YTHDF2 as a negative regulator for dendrite development and maintenance of retinal ganglion cells (RGCs) ***(Niu et al. 2022)***, which inspired us to further explore whether YTHDF2 regulates RGC dendrite morphonology in the aged mice.

We kept *Six3-cre*^*+/-*^,*Ythdf2*^*fl/fl*^ (*Ythdf2* cKO) and *Ythdf2*^*fl/fl*^ control mice and had them grow to 23-25 months old, which is equivalent to approximately 70 years old for human. We first checked the dendrite morphology of ipRGCs with anti-melanopsin immunostaining in early adult and aged mouse retinas. The dendritic area of melanopsin^+^ ipRGCs was significantly decreased in the aged control mice (23∼25 months old) compared with young adult control mice (3.5 months old) (***Figure 1A***), which is consistent with the previous studies ***(Samuel et al. 2011)***. Next, we continued to examine the dendrite morphology of ipRGCs in the aged *Ythdf2* cKO and control mice. The dendritic area of ipRGCs was significantly large in the aged *Ythdf2* cKO mice (***Figure 1B and C***). The ipRGCs of the aged *Ythdf2* cKO mice exhibited more dendrite branches and higher complexity compared to the age-matched controls (***Figure 1B and D***). All these results suggest that the shrinking of dendrite area and complexity of ipRGCs caused by aging is dramatically alleviated in the aged *Ythdf2* cKO mice (***Figure 1B-D***).

**Figure 1.**
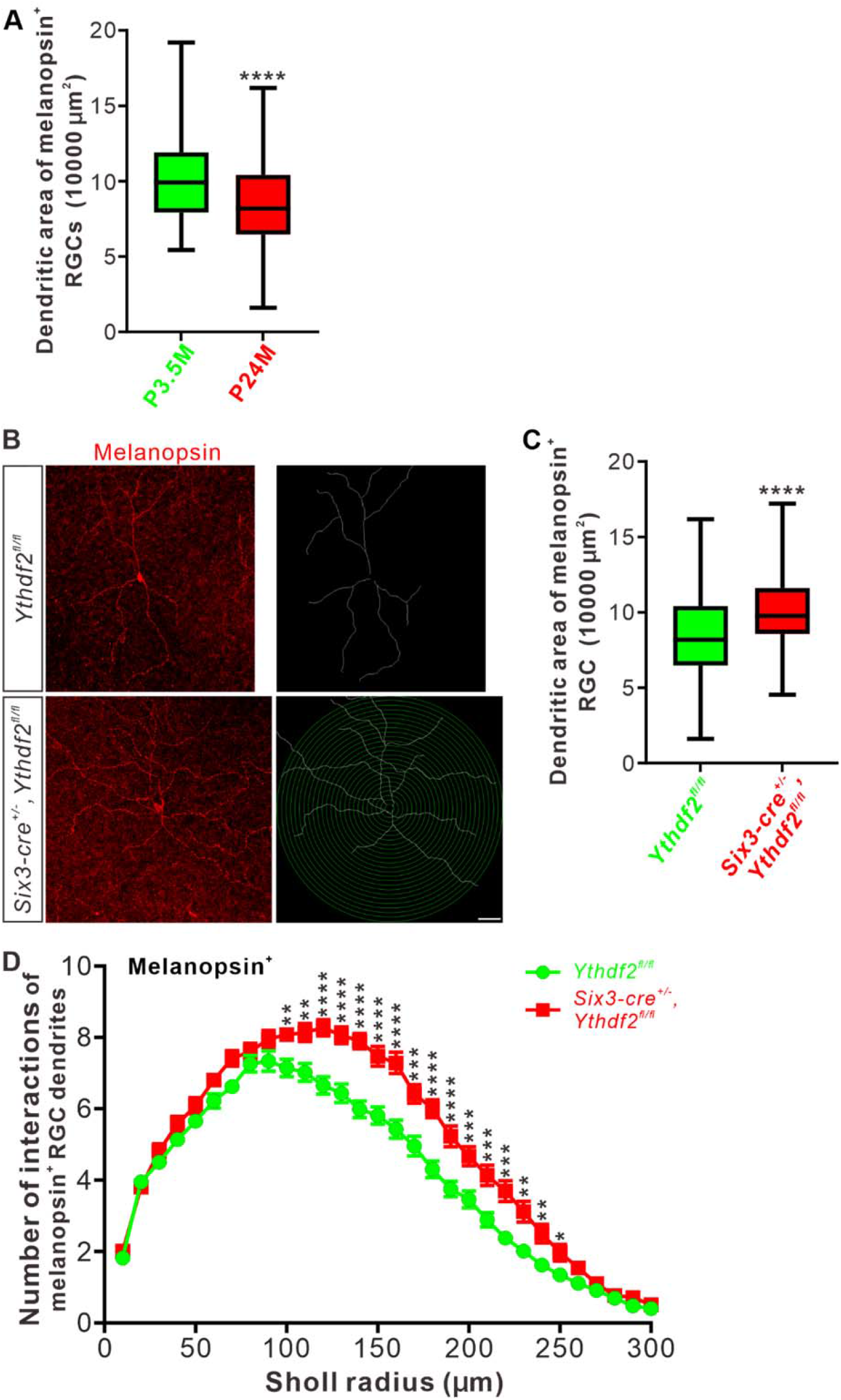
*Ythdf2* cKO protects retina from age-related RGC dendrite shrinking. **A**. Quantification of dendritic area of melanopsin^+^ ipRGCs in young adult and aged mice. Data are represented as box and whisker plots: **** *p* = 1.33E-05 (*n* = 71 RGCs for P3.5M, n = 89 B.RGCs for P24M); by unpaired Student’s *t*-test. P3.5M, postnatal 3.5 months; P24M, postnatal 24 months. **B**. Representative images of wholemount immunostaining of 23-25 months old mouse retina using a melanopsin antibody. Scale bar: 50 μm. **C**. Quantification of dendritic areas of melanopsin^+^ ipRGCs (**B**). Aged *Ythdf2* cKO mice maintain significantly larger dendritic areas compared with age-matched control mice: \*\*\*\**p* = 1.62E-05 (*n* = 89 RGCs for control, n = 67 RGCs for cKO); by unpaired Student’s *t*-test. **D**. Quantification of dendrite branching of melanopsin^+^ ipRGCs (**B**) using Sholl analysis. Data are mean ± SEM. Numbers of interactions are significantly greater in *Six3-cre*^*+/-*^,*Ythdf2*^*fl/fl*^ groups (*n* = 66 RGCs) than *Ythdf2*^*fl/fl*^ groups (*n* = 89 RGCs) in Sholl radii between 100-250 μm: \*\**p* = 0.0051 (100 μm), \*\**p* = 0.0025 (110 μm), \*\*\*\**p* = 6.24E-06 (120 μm), \*\*\*\**p* = 1.37E-05 (130 μm), \*\*\*\**p* = 5.21E-08 (140 μm), \*\*\*\**p* = 1.81E-05 (150 μm), \*\*\*\**p* = 4.61E-06 (160 μm), \*\*\**p* = 0.00022 (170 μm), \*\*\*\**p* = 3.87E-06 (180 μm), \*\*\*\**p* = 7.60E-05 (190 μm), \*\*\**p* = 0.00084 (200 μm), \*\*\**p* = 0.00031 (210 μm), \*\*\**p* = 0.00011 (220 μm), \*\**p* = 0.0010 (230 μm), \*\**p* = 0.0037 (240 μm), \**p* = 0.023 (250 μm), by unpaired Student’s *t*-test.

### Visual acuity is improved in the aged *Ythdf2* cKO mice

The aging-related declines in spatial contrast sensitivity and visual acuity are attributed to neuronal changes, such as degeneration of RGC dendrite or axons ***(Samuel et al. 2011)***. The better RGC dendrite maintenance in the aged *Ythdf2* cKO mice inspired us to further explore whether the visual responses of the aged *Ythdf2* cKO mice were improved or not.

The aged *Ythdf2* cKO mice showed similar body size and weight as controls (***Figure 2A and B***). We carried out the optomotor test on these aged mice. The aged control mice revealed significantly decreased visual acuity with the spatial frequency threshold as 0.32 ± c/deg (***Figure 2C***), compared with the young adult control mice measuring 0.43 ± 0.0085 c/deg ***(Niu et al. 2022)***. However, the aged *Ythdf2* cKO mice show significantly better visual acuity (0.38 ± 0.011 c/deg) compared with the aged control (***Figure 2C***). These data suggest that the ablation of *Ythdf2* in retina improves visual acuity of the aged mice.

**Figure 2.**
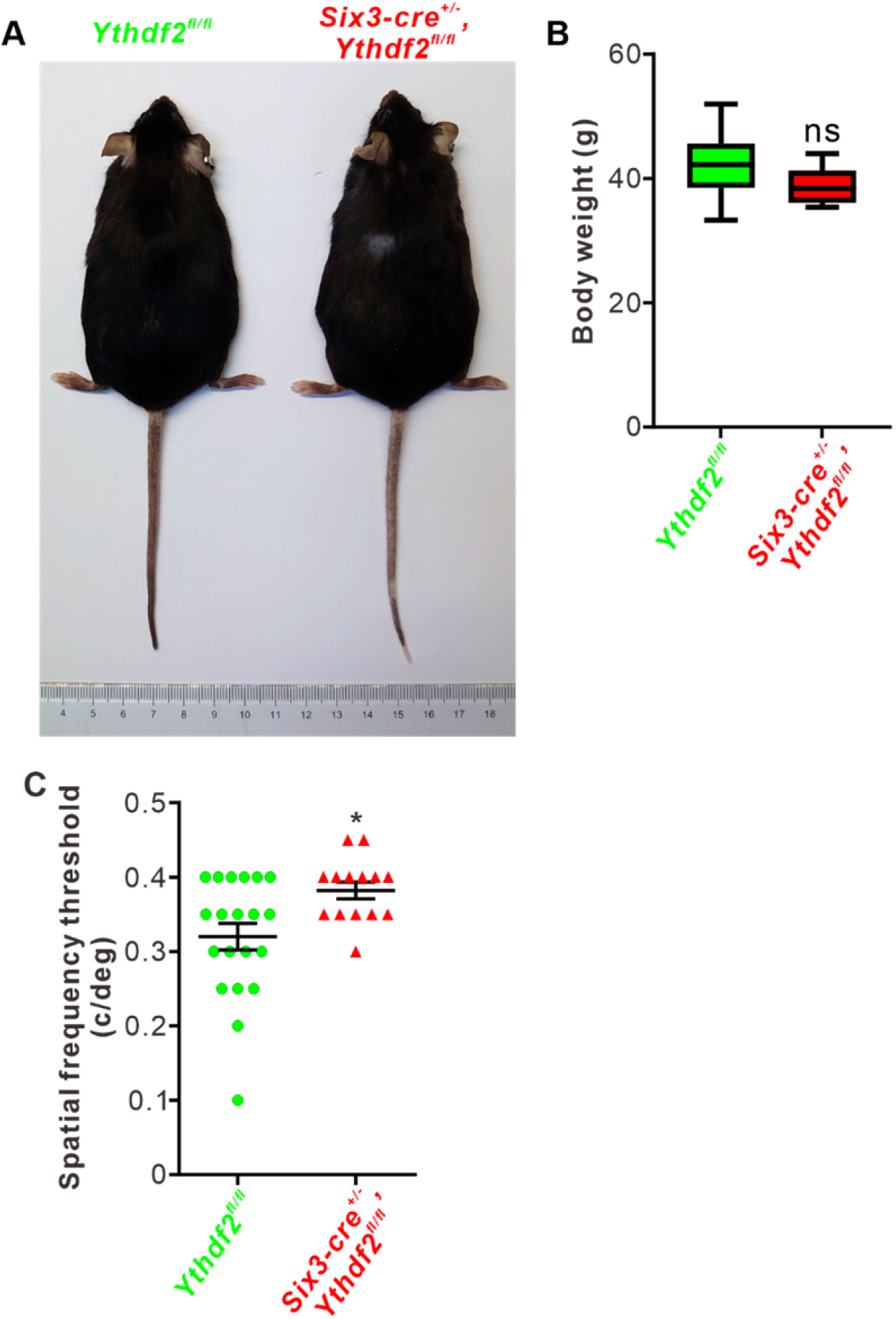
Visual acuity is improved in the aged *Ythdf2* cKO mice. **A**. The aged *Ythdf2* cKO mice show similar body size as the control mice. **B**. Quantification of body weight of the aged *Ythdf2* cKO and control mice. Data are represented as box and whisker plots: *p* = 0.18 (*n* = 10 for control, *n* = 7 for cKO); ns, not significant; by unpaired Student’s *t*-test. **C**. The aged *Ythdf2* cKO mice demonstrate better visual acuity. Quantification data are mean ± SEM: \**p* = 0.012 (*n* = 20 control, *n* = 14 cKO; all male); by unpaired Student’s *t*-test.

### *Hspa12a* and *Islr2* are the YTHDF2 target mRNAs in the aged RGCs

By proteomic analysis and anti-YTHDF2 RNA immunoprecipitation sequencing, we have identified *Hspa12a* and *Islr2* as the YTHDF2 targets in a glaucoma model ***(Niu et al. 2022)***. We further investigated whether *Hspa12a* and *Islr2* mediate RGC dendrite shrinking in the aged retinas. We firstly performed RT-qRCR to check the expression levels of *Hspa12a* and *Islr2* by comparing the aged *Ythdf2* cKO and control retinas. Upregulation of their expression levels was detected in the aged *Ythdf2* cKO retina (***Figure 3A***), implying that the improved visual function in the *Ythdf2* cKO mice is likely mediated by the neuroprotective YTHDF2 targets *Hspa12a* and *Islr2*.

**Figure 3.**
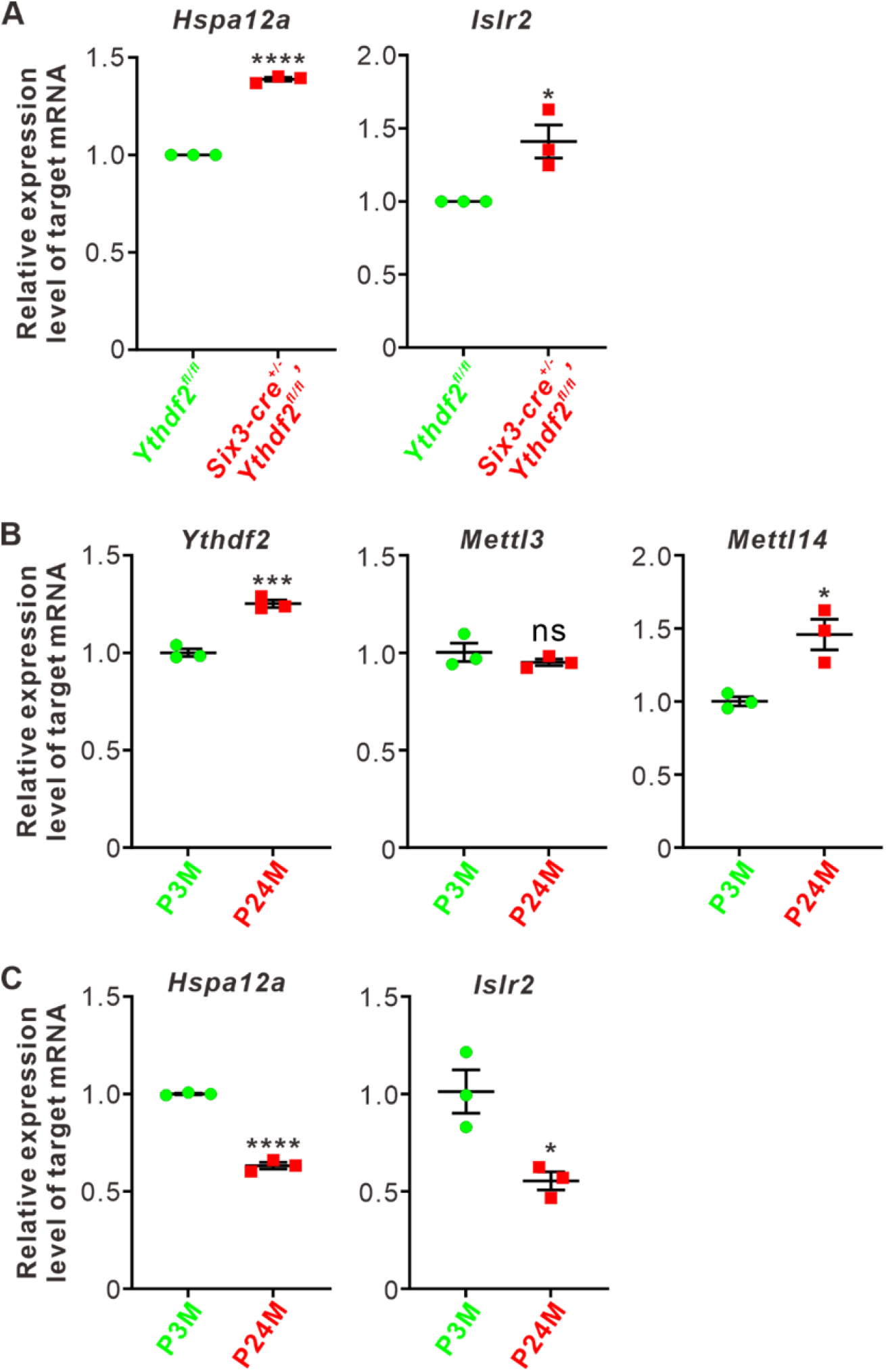
*Hspa12a* and *Islr2* are the YTHDF2 targets in the aged retinas. **A**. Upregulation of YTHDF2 target mRNAs *Hspa12a* and *Islr2* in P24M *Ythdf2* cKO retina compared with control by RT-qPCR. Data are mean ± SEM and are represented as dot plots (*n* = 3 replicates): \*\*\*\**p* = 2.97E-05 for *Hspa12a*; \**p* = 0.023 for *Islr2*; by unpaired Student’s *t*-test. **B**.Upregulation of *Ythdf2* and *Mettl14* mRNA levels in the aged retinas. Data are mean ± SEM and are represented as dot plots (*n* = 3 replicates): \*\*\**p* = 0.00081 for *Ythdf2, p* = 0.37 for *Mettl3*, \**p* = 0.014 for *Mettl14*; ns, not significant; by unpaired Student’s *t*-test. **C**. Downregulation of *Hspa12a* and *Islr2* mRNA levels in the aged retinas. Data are mean ± SEM and are represented as dot plots (*n* = 3 replicates): \*\*\*\**p* = 2.63E-05 for *Hspa12a*; \**p* = for *Islr2*; by unpaired Student’s *t*-test.

Next, we continued to explore this pathway in the normal aging progress. We found that expression of *Mettl14* and *Ythdf2* was upregulated in the aged mouse retina compared with the young adults, although *Mettl3* expression was not changed (***Figure 3B***). In line with this, *Hspa12a* and *Islr2* mRNA levels were downregulated with aging (***Figure 3C***). The upregulation of the m^6^A writer *Mettl14* and reader *Ythdf2* in the aged retinas might account for the downregulation of *Hspa12a* and *Islr2*.

Together, these data suggest that *Ythdf2* cKO protects the aged retina from aging-related RGC dendrite shrinking and visual loss, possibly through avoiding downregulation of its neuroprotective targets, *Hspa12a* and *Islr2*.

## Discussion

In the previous work, we revealed that YTHDF2 is a negative regulator for dendrite development and maintenance of RGCs. Loss-of function of YTHDF2 induces increased RGC dendrite branching, more synapses, improved visual acuity, and more resistant for glaucoma model ***(Niu et al. 2022)***. In this study, we further explored the function of YTHDF2 to mediate m^6^A modification in RGC degeneration and vision loss by aging. In the aged retinas, *Ythdf2* ablation disrupts the de-stabilization of its target mRNAs and thus increases the levels of its target mRNAs including *Hspa12a*, and *Islr2*, which results in less RGC dendrite shrinking and less visual loss (***Figure 4***).

**Figure 4.**
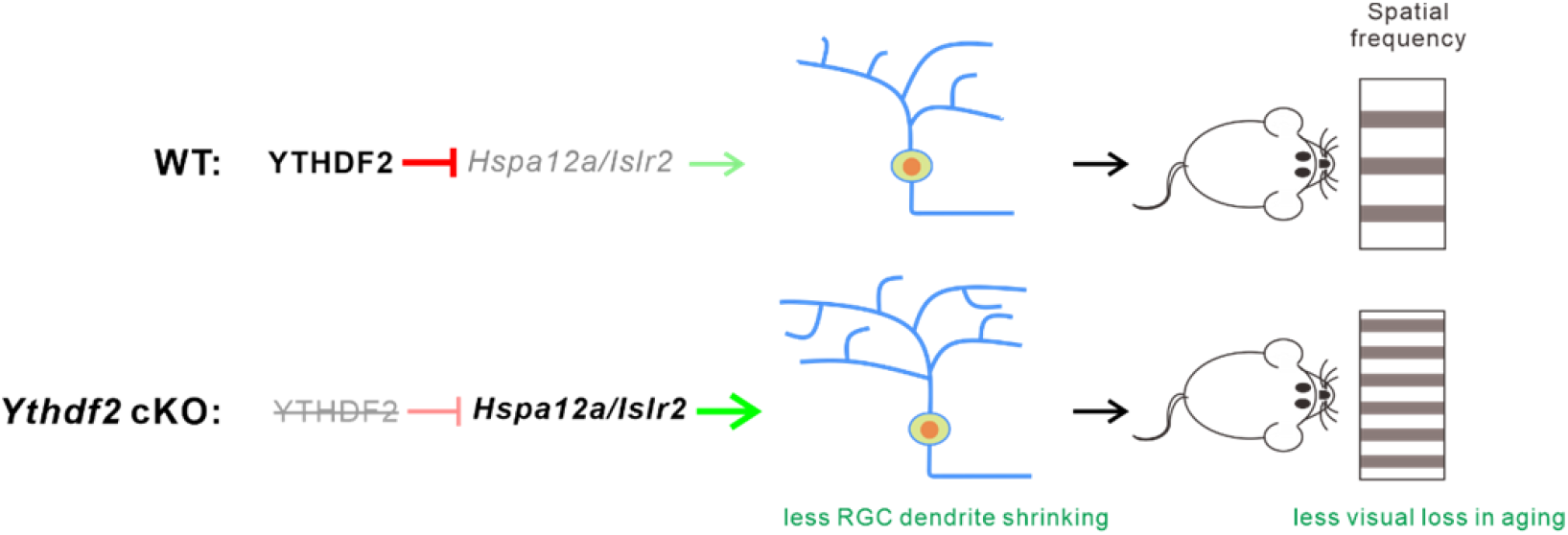
A proposed model for how YTHDF2 regulates aging-related neurodegeneration in retina through its target mRNAs. Normally YTHDF2 downregulates its target mRNAs *Hspa12a* and *Islr2* in the aged retinas, which leads to RGC dendrite shrinking and vision loss. Ablation of *Ythdf2* increases *Hspa12a* and *Islr2* levels in the aged retinas, which results in less RGC dendrite shrinking and less visual loss with aging.

The precise mechanisms of how the aging upregulates m^6^A modification and how the YTHDF2 target mRNAs protect neurodegeneration in retina still requires further investigation. Nevertheless, the epitranscriptomic regulation through m^6^A modification in gene expression can be evaluated as possible therapeutic targets for aging-related vision decline.

## Materials and methods

### Key resources table

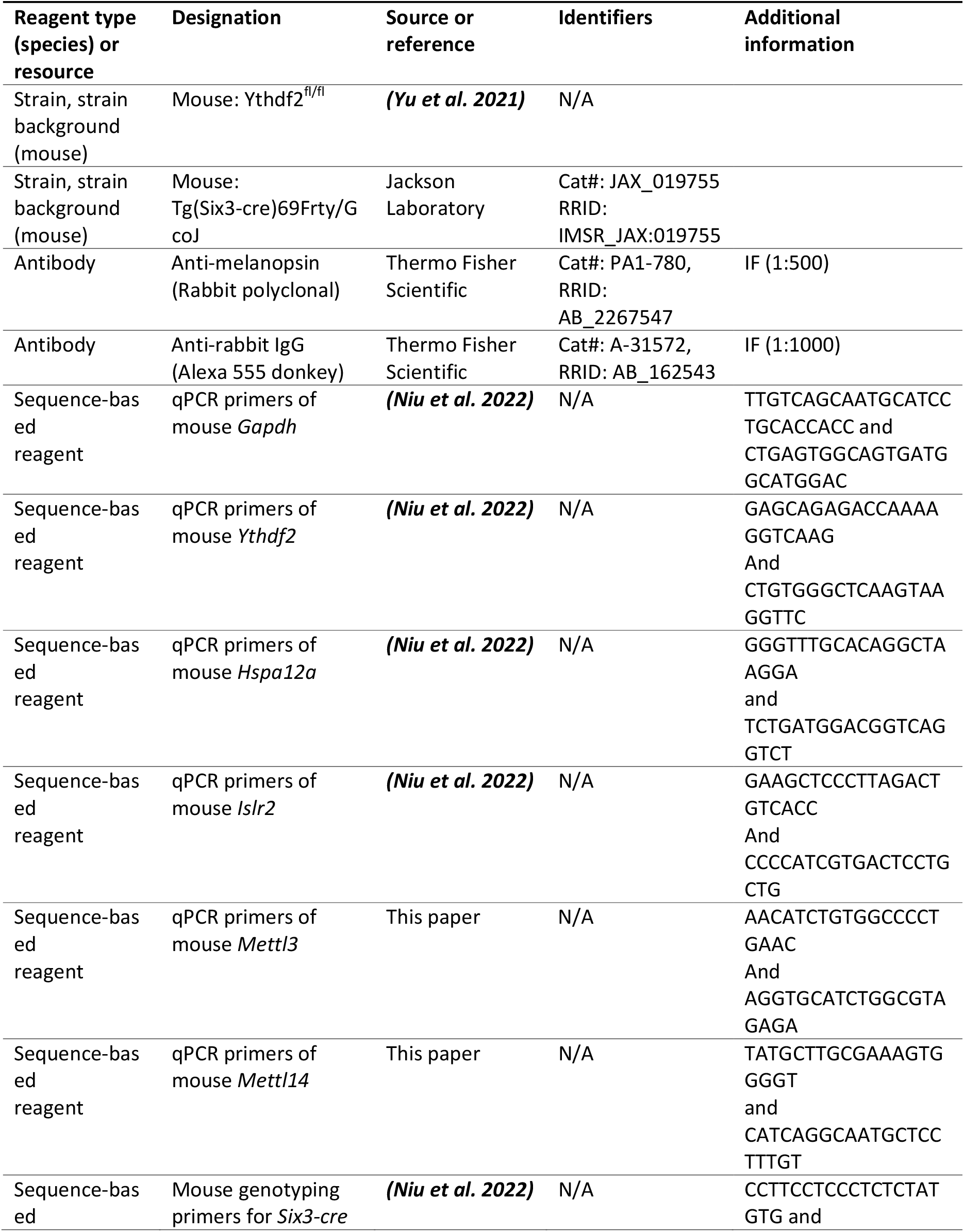
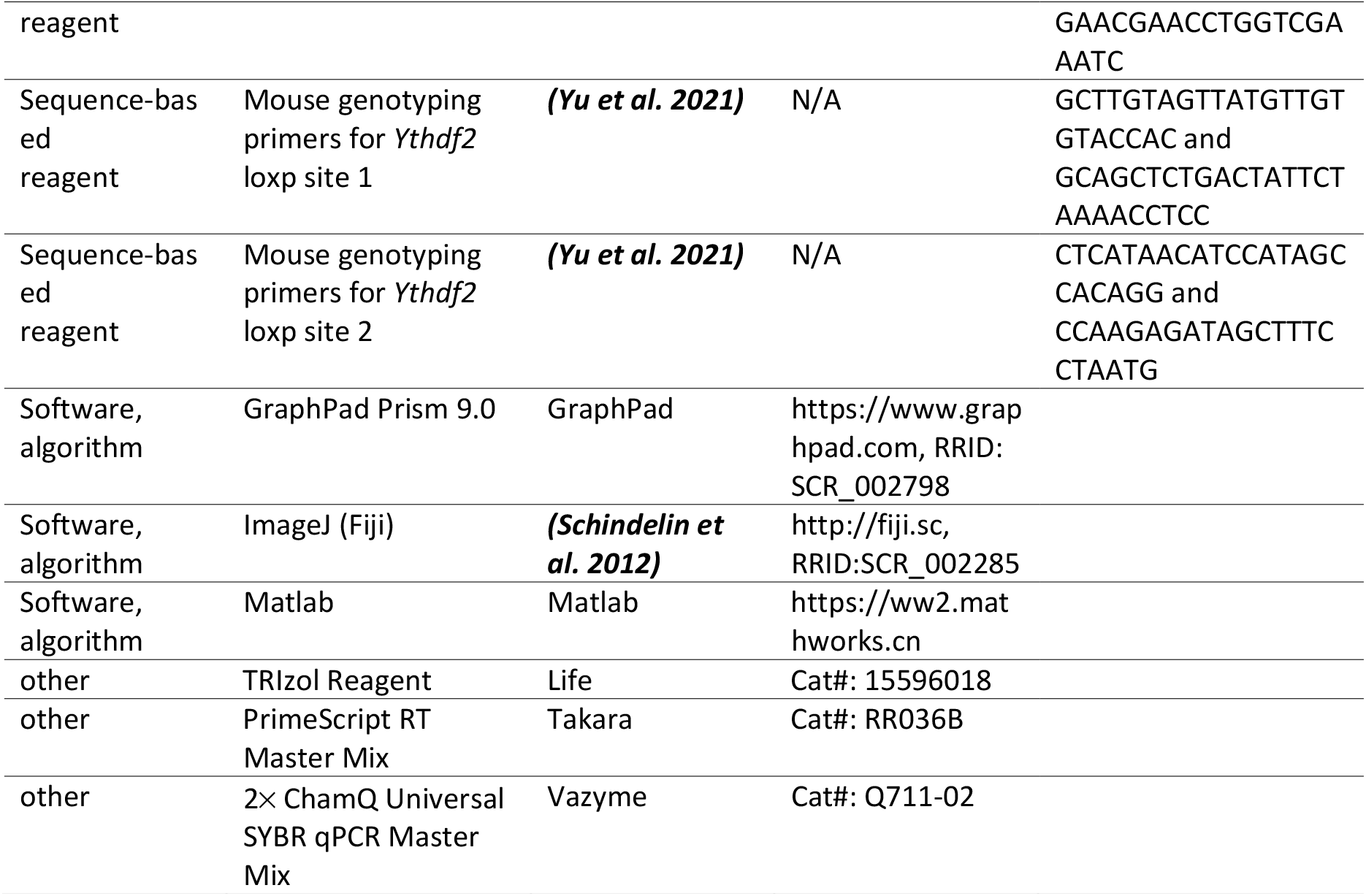

### Animals

*Ythdf2*^*fl/fl*^ mice were reported previously ***(Yu et al. 2021)***, and *Six3-cre* ***(Furuta et al. 2000)*** (The Jackson Laboratory, # 019755) were from Jackson Laboratory. Genotyping primers are as follows: The first *Ythdf2*-loxp site: 5’-GCTTGTAGTTATGTTGTGTACCAC-3’ and 5’-GCAGCTCTGACTATTCTAAAACCTCC-3’; the second *Ythdf2*-loxp site: 5’-CTCATAACATCCATAGCCACAGG-3’ and 5’-CCAAGAGATAGCTTTCCTAATG-3’. *Six3-cre* site: 5’-CCTTCCTCCCTCTCTATGTG-3’ and 5’-GAACGAACCTGGTCGAAATC-3’. All experiments using mice were carried out following the animal protocols approved by the Laboratory Animal Welfare and Ethics Committee of Southern University of Science and Technology.

### Immunostaining

For anti-melanopsin retinal wholemount staining, the process was described previously ***(Niu et al. 2022)***. All images were captured on Zeiss LSM 800 confocal microscope with identical settings for each group in the same experiment.

### RT-qPCR

Total RNA was extracted from retinas with TRIzol Reagent (Life) and then used for reverse transcription by PrimeScript RT Master Mix (TaKaRa). Synthesized cDNA was performed with 2× ChamQ Universal SYBR qPCR Master Mix (Vazyme) on BioRad CFX96 Touch Real-Time PCR system. Primers used for qPCR are as following: mouse *Gapdh*:

5’-TTGTCAGCAATGCATCCTGCACCACC-3’ and 5’-CTGAGTGGCAGTGATGGCATGGAC-3’ ***(Niu et al. 2022)***; mouse *Ythdf2*: 5’-GAGCAGAGACCAAAAGGTCAAG-3’and 5’-CTGTGGGCTCAAGTAAGGTTC-3’ ***(Niu et al. 2022)***; mouse *Hspa12a*:

5’-GGGTTTGCACAGGCTAAGGA-3’ and 5’-TCTGATGGACGGTCAGGTCT-3’ ***(Niu et al. 2022)***; mouse *Islr2*: 5’-GAAGCTCCCTTAGACTGTCACC-3’ and 5’-CCCCATCGTGACTCCTGCTG-3’ ***(Niu et al. 2022)***. Mouse *Mettl3*: 5’-AACATCTGTGGCCCCTGAAC-3’ and 5’-AGGTGCATCTGGCGTAGAGA-3’; mouse *Mettl14*: 5’-TATGCTTGCGAAAGTGGGGT-3’ and 5’-CATCAGGCAATGCTCCTTTGT-3’.

### Optomotor response (OMR) assay

*Ythdf2* cKO and control mice aged about 24 months were applied for OMR assay as previously reported ***(Niu et al. 2022)***. Using the Matlab program, 0.075, 0.1, 0.2, 0.25, 0.3, 0.35, 0.4, 0.45 and 0.5 c/deg (30s per direction of rotation) were used in the recording process. Mouse behaviors were analyzed in real time during the experiment and re-checked with the video recordings. Finally, the minimal spatial frequency of left and right OMR for each mouse was recorded and analyzed respectively.

### Quantification and statistical analysis

All experiments were conducted at a minimum of three independent biological replicates in the lab. Statistical analysis was performed using GraphPad Prism 9.0. When comparing the means of two groups, an unpaired *t*-test was performed on the basis of experimental design. The settings for all box and whisker plots are: 25th-75th percentiles (boxes), minimum and maximum (whiskers), and medians (horizontal lines). Data for all other graphs are mean ± SEM. A *p* value less than 0.05 was considered as statistically significant: \**p* < 0.05, \*\**p* < 0.01, \*\*\**p* < 0.001, \*\*\*\**p* < 0.0001.

## Acknowledgements

We thank other members of Ji laboratory for technical support, helpful discussions and comments on the manuscript. We thank the technical support from the Laboratory Animal Center and the Core Research Facilities of Southern University of Science and Technology. This work was supported by National Natural Science Foundation of China (31871038 and 32170955 to S.-J.J.), Shenzhen-Hong Kong Institute of Brain Science-Shenzhen Fundamental Research Institutions (2022SHIBS0002), High-Level University Construction Fund for Department of Biology (internal grant no. G02226301), Science and Technology Innovation Commission of Shenzhen Municipal Government (ZDSYS20200811144002008).

## Ethics

All experiments using mice were carried out following the animal protocols approved by the Laboratory Animal Welfare and Ethics Committee of Southern University of Science and Technology (approval numbers: SUSTC-JY2017004, SUSTC-JY2019081).

## Data availability

This work did not generate any dataset. Figure 1 - Source Data 1, Figure 2 - Source Data 1, and Figure 3 - Source Data 1 contain the numerical data used to generate the figures.

## References

Blindness GBD, Vision Impairment C, Vision Loss Expert Group of the Global Burden of Disease S. 2021. Causes of blindness and vision impairment in 2020 and trends over 30 years, and prevalence of avoidable blindness in relation to VISION 2020: the Right to Sight: an analysis for the Global Burden of Disease Study. The Lancet. Global health 9:e144–e160. DOI: https://doi.org/10.1016/S2214-109X(20)30489-7, PMID: 33275949

Datta S, Cano M, Ebrahimi K, Wang L, Handa JT. 2017. The impact of oxidative stress and inflammation on RPE degeneration in non-neovascular AMD. Prog Retin Eye Res 60:201–218. DOI: https://doi.org/10.1016/j.preteyeres.2017.03.002, PMID: 28336424

Esquiva G, Lax P, Pérez-Santonja JJ, García-Fernández JM, Cuenca N. 2017. Loss of Melanopsin-Expressing Ganglion Cell Subtypes and Dendritic Degeneration in the Aging Human Retina. Frontiers in Aging Neuroscience 9. DOI: https://doi.org/10.3389/fnagi.2017.00079,

Flaxman SR, Bourne RRA, Resnikoff S, Ackland P, Braithwaite T, Cicinelli MV, Das A, Jonas JB, Keeffe J, Kempen JH, Leasher J, Limburg H, Naidoo K, Pesudovs K, Silvester A, Stevens GA, Tahhan N, Wong TY, Taylor HR. 2017. Global causes of blindness and distance vision impairment 1990-2020: a systematic review and meta-analysis. Lancet Glob Health 5:e1221–e1234. DOI: https://doi.org/10.1016/s2214-109x(17)30393-5, PMID: 29032195

Fleckenstein M, Keenan TDL, Guymer RH, Chakravarthy U, Schmitz-Valckenberg S, Klaver CC, Wong WT, Chew EY. 2021. Age-related macular degeneration. Nat Rev Dis Primers 7:31. DOI: https://doi.org/10.1038/s41572-021-00265-2, PMID: 33958600

Furuta Y, Lagutin O, Hogan BL, Oliver GC. 2000. Retina- and ventral forebrain-specific Cre recombinase activity in transgenic mice. Genesis 26:130-132. PMID: 10686607

Jackson GR, Owsley C, Curcio CA. 2002. Photoreceptor degeneration and dysfunction in aging and age-related maculopathy. Ageing Res Rev 1:381–396. DOI: https://doi.org/10.1016/s1568-1637(02)00007-7, PMID: 12067593

Koch SC, Nelson A, Hartenstein V. 2021. Structural aspects of the aging invertebrate brain. Cell Tissue Res 383:931–947. DOI: https://doi.org/10.1007/s00441-020-03314-6, PMID: 33409654

Lefebvre JL, Sanes JR, Kay JN. 2015. Development of dendritic form and function. Annu Rev Cell Dev Biol 31:741–777. DOI: https://doi.org/10.1146/annurev-cellbio-100913-013020, PMID: 26422333

Masland RH. 2012. The neuronal organization of the retina. Neuron 76:266–280. DOI: https://doi.org/10.1016/j.neuron.2012.10.002, PMID: 23083731

Niu F, Han P, Zhang J, She Y, Yang L, Yu J, Zhuang M, Tang K, Shi Y, Yang B, Liu C, Peng B, Ji SJ. 2022. The m(6)A reader YTHDF2 is a negative regulator for dendrite development and maintenance of retinal ganglion cells. Elife 11. DOI: https://doi.org/10.7554/eLife.75827, PMID: 35179492

Owsley C. 2016. Vision and Aging. Annu Rev Vis Sci 2:255–271. DOI: https://doi.org/10.1146/annurev-vision-111815-114550, PMID: 28532355

Pelletier AL, Rojas-Roldan L, Coffin J. 2016. Vision Loss in Older Adults. Am Fam Physician 94:219-226. PMID: 27479624

Qu X, Zhu K, Li Z, Zhang D, Hou L. 2021. The Alteration of M6A-Tagged Transcript Profiles in the Retina of Rats After Traumatic Optic Neuropathy. Front Genet 12:628841. DOI: https://doi.org/10.3389/fgene.2021.628841, PMID: 33664770

Samuel MA, Zhang Y, Meister M, Sanes JR. 2011. Age-related alterations in neurons of the mouse retina. J Neurosci 31:16033–16044. DOI: https://doi.org/10.1523/jneurosci.3580-11.2011, PMID: 22049445

Schindelin J, Arganda-Carreras I, Frise E, Kaynig V, Longair M, Pietzsch T, Preibisch S, Rueden C, Saalfeld S, Schmid B, Tinevez JY, White DJ, Hartenstein V, Eliceiri K, Tomancak P, Cardona A. 2012. Fiji: an open-source platform for biological-image analysis. Nat Methods 9:676–682. DOI: https://doi.org/10.1038/nmeth.2019, PMID: 22743772

Spear PD. 1993. Neural bases of visual deficits during aging. Vision Res 33:2589–2609. DOI: https://doi.org/10.1016/0042-6989(93)90218-l, PMID: 8296455

Suo L, Liu C, Zhang QY, Yao MD, Ma Y, Yao J, Jiang Q, Yan B. 2022. METTL3-mediated N (6)-methyladenosine modification governs pericyte dysfunction during diabetes-induced retinal vascular complication. Theranostics 12:277–289. DOI: https://doi.org/10.7150/thno.63441, PMID: 34987645

Weinreb RN, Aung T, Medeiros FA. 2014. The Pathophysiology and Treatment of Glaucoma: A Review. JAMA 311:1901–1911. DOI: https://doi.org/10.1001/jama.2014.3192,

Yao MD, Jiang Q, Ma Y, Liu C, Zhu CY, Sun YN, Shan K, Ge HM, Zhang QY, Zhang HY, Yao J, Li XM, Yan B. 2020. Role of METTL3-Dependent N(6)-Methyladenosine mRNA Modification in the Promotion of Angiogenesis. Mol Ther 28:2191–2202. DOI: https://doi.org/10.1016/j.ymthe.2020.07.022, PMID: 32755566

Yu J, She Y, Yang L, Zhuang M, Han P, Liu J, Lin X, Wang N, Chen M, Jiang C, Zhang Y, Yuan Y, Ji SJ. 2021. The m(6) A Readers YTHDF1 and YTHDF2 Synergistically Control Cerebellar Parallel Fiber Growth by Regulating Local Translation of the Key Wnt5a Signaling Components in Axons. Adv Sci (Weinh) 8:e2101329. DOI: https://doi.org/10.1002/advs.202101329, PMID: 34643063

